# A bacterial extracellular vesicle-based intranasal vaccine against SARS-CoV-2 protects against disease and elicits neutralizing antibodies to wild-type and Delta variants

**DOI:** 10.1101/2021.06.28.450181

**Authors:** Linglei Jiang, Tom A.P. Driedonks, Wouter S.P. Jong, Santosh Dhakal, H. Bart van den Berg van Saparoea, Ioannis Sitaras, Ruifeng Zhou, Christopher Caputo, Kirsten Littlefield, Maggie Lowman, Mengfei Chen, Gabriela Lima, Olesia Gololobova, Barbara Smith, Vasiliki Mahairaki, M. Riley Richardson, Kathleen R. Mulka, Andrew P. Lane, Sabra L. Klein, Andrew Pekosz, Cory F. Brayton, Joseph L. Mankowski, Joen Luirink, Jason S. Villano, Kenneth W. Witwer

## Abstract

Several vaccines have been introduced to combat the coronavirus infectious disease-2019 (COVID-19) pandemic, caused by severe acute respiratory syndrome coronavirus 2 (SARS-CoV-2). Current SARS-CoV-2 vaccines include mRNA-containing lipid nanoparticles or adenoviral vectors that encode the SARS-CoV-2 Spike (S) protein of SARS-CoV-2, inactivated virus, or protein subunits. Despite growing success in worldwide vaccination efforts, additional capabilities may be needed in the future to address issues such as stability and storage requirements, need for vaccine boosters, desirability of different routes of administration, and emergence of SARS-CoV-2 variants such as the Delta variant. Here, we present a novel, well-characterized SARS-CoV-2 vaccine candidate based on extracellular vesicles (EVs) of *Salmonella typhimurium* that are decorated with the mammalian cell culture-derived Spike receptor-binding domain (RBD). RBD-conjugated outer membrane vesicles (RBD-OMVs) were used to immunize the golden Syrian hamster (*Mesocricetus auratus*) model of COVID-19. Intranasal immunization resulted in high titers of blood anti-RBD IgG as well as detectable mucosal responses. Neutralizing antibody activity against wild-type and Delta variants was evident in all vaccinated subjects. Upon challenge with live virus, hamsters immunized with RBD-OMV, but not animals immunized with unconjugated OMVs or a vehicle control, avoided body mass loss, had lower virus titers in bronchoalveolar lavage fluid, and experienced less severe lung pathology. Our results emphasize the value and versatility of OMV-based vaccine approaches.

## INTRODUCTION

The coronavirus infectious disease-2019 (COVID-19) pandemic, caused by severe acute respiratory syndrome coronavirus 2 (SARS-CoV-2) ^1,2^, has highlighted the need for rapid vaccine development capabilities ^3^. Current SARS-CoV-2 vaccines consist of mRNA-containing lipid nanoparticles or adenoviral vectors that encode the surface Spike (S) protein of SARS-CoV-2 ^4–6^. Other vaccination approaches use inactivated virus or protein subunits ^7^. Several of the available vaccines have elicited remarkable protection against disease ^8,9^, and worldwide vaccination efforts have achieved tremendous successes in many countries. Despite this progress, factors such as stability and storage requirements, speed of reaction, and production scalability may make novel approaches desirable to combat new variants of SARS-CoV-2 or future emerging viruses. SARS-CoV-2 has accumulated mutations during the COVID-19 pandemic, and a subset of lineages have been designated as variants of concern (VOC) due to increased transmission, escape from vaccine-induced immunity, or morbidity and mortality. Recently, the B.1.6.17.2 (Delta) variant has become the dominant lineage in several countries, is reported to be more transmissible than previously found variants, and evades some of the antibody responses induced in humans vaccinated with the vaccines including the Pfizer and Moderna vaccines ^10,11^.

Here, we present a novel SARS-CoV-2 vaccine candidate based on bacterial extracellular vesicles (EVs) that are decorated with the Spike receptor-binding domain (RBD). Gram-negative bacteria such as *Salmonella typhimurium* produce EVs known as outer membrane vesicles (OMVs). These vesicles, like their parent cells, have endotoxin-mediated immunostimulatory properties in mammalian hosts, driving inflammation and potently activating immune cells including dendritic cells, T cells, and B cells ^12,13^. Although native bacterial OMVs can elicit damaging systemic responses ^14^, OMVs can also be prepared from engineered, endotoxin-attenuated bacteria ^15^. We prepared OMVs from an attenuated strain of *S. typhimurium* displaying a version of the virulence factor hemoglobin protease (Hbp) that carries the SpyCatcher peptide for coupling of protein cargo containing a SpyTag ^16^. The SpyTag/SpyCatcher system enables coupling of proteins *via* a covalent amide bond that is stable under broad pH, temperature and buffer conditions ^17^. We report that this technology efficiently couples a SpyTag-RBD fusion protein produced in mammalian cell culture onto bacterial OMVs, resulting in RBD-OMVs that are recognized by antibodies against SARS-CoV-2. Furthermore, we show that intranasal vaccination with RBD-OMVs elicits antibodies, including neutralization responses against both wild-type and Delta viral variants, and confers protection against challenge with SARS-CoV-2 in a recently developed hamster model ^18,19^.

## RESULTS

We designed expression constructs to produce RBD domain of SARS-CoV2-Spike harboring SpyTag and 6xHis-tag motifs on the N-terminal or C-terminal end (Figure 1A). This allows coupling of RBD to OMVs from detoxified *S. typhimurium* displaying Hbp modified with the SpyCatcher peptide (Figure 1B).

**Figure 1.**
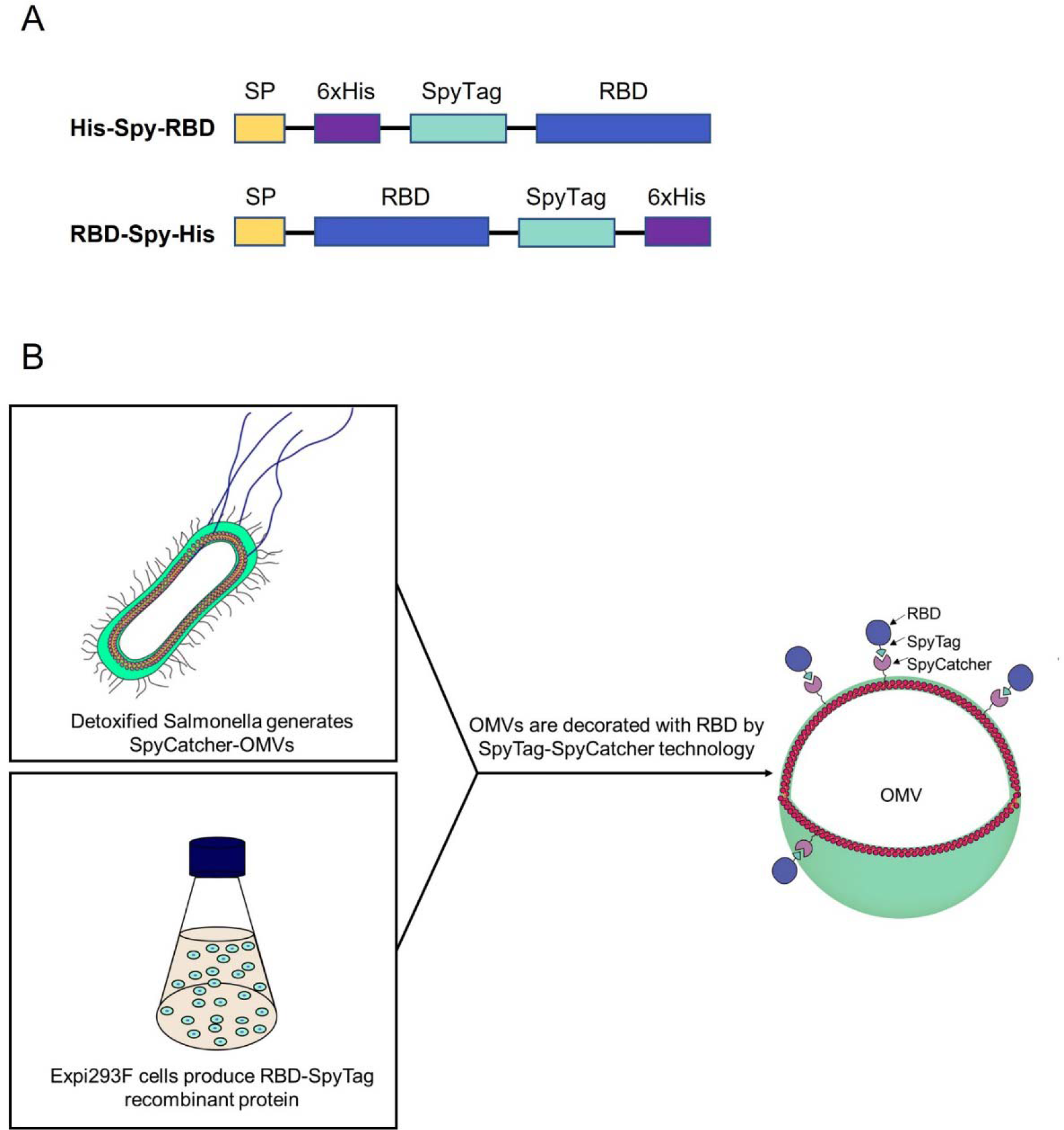
Schematic of expression constructs and OMV decoration. A) Design of RBD recombinant antigens fused to N- and C-terminal SpyTag. B) Schematic representation of the production of RBD-OMVs.

Efficient coupling of RBD-Spy-His and Spy-His-RBD to HbpD was demonstrated by SDS-PAGE and Coomassie staining, showing that virtually all of the exposed HbpD was coupled to RBD independent of the orientation of SpyTag (Figure 2A). OMV batches carrying RBD with either N-or C-terminal SpyTag were blended in a 1:1 ratio to produce a vaccine formulation (RBD-OMV), whereas native, non-conjugated OMVs were used as a control (Ctrl-OMV) (Figure 2B). The N-glycosylation state of RBD was confirmed by immunoblotting with/without prior PNGase F treatment. (Supplementary Figure S1A). Successful decoration of RBD onto the surface of OMVs was further confirmed by Western blot. Lipopolysaccharide (LPS), as expected, was associated with both RBD-OMV and Ctrl-OMV (Supplementary Figure S1B). Detection of RBD with anti-His and anti-Spike antibodies showed specific bands with the expected molecular weight of approximately 160 kDa (Figure 3B and Supplementary Figure S1C).

**Figure 2.**
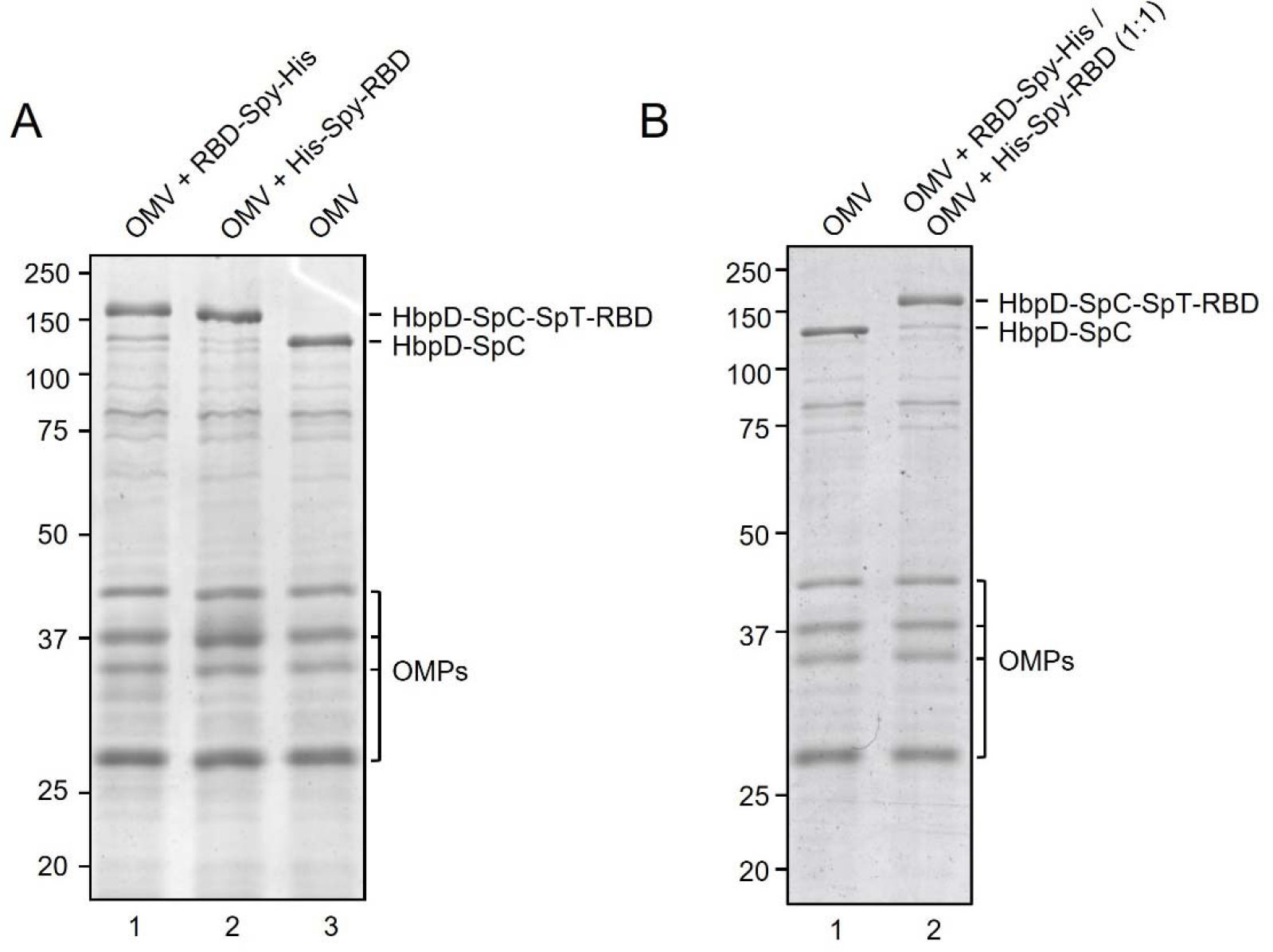
**A)** Assessment of efficiency of SpyTag/SpyCatcher coupling of RBD onto HbpD of OMVs. RBD-Spy-His and His-Spy-RBD were coupled to Hbp-SpyCatcher OMVs. Proteins of conjugated and non-conjugated OMVs were separated by SDS-PAGE and stained with Coomassie Brilliant Blue. RBD-HbpD appears as a ∼160 kDa band, while free HbpD is seen as a ∼125 kDa band. Densitometry suggested that approximately 90% or more of HbpD was coupled with RBD in the conjugated populations compared with unconjugated OMVs (rightmost lane). Other outer membrane proteins of OMVs (OMPs) are indicated; B) Coomassie Brilliant Blue staining of SDS-PAGE gel containing non-conjugated OMVs and a 1:1 mixture of RBD-Spy-His and His-Spy-RBD-coupled OMVs.

**Figure 3.**
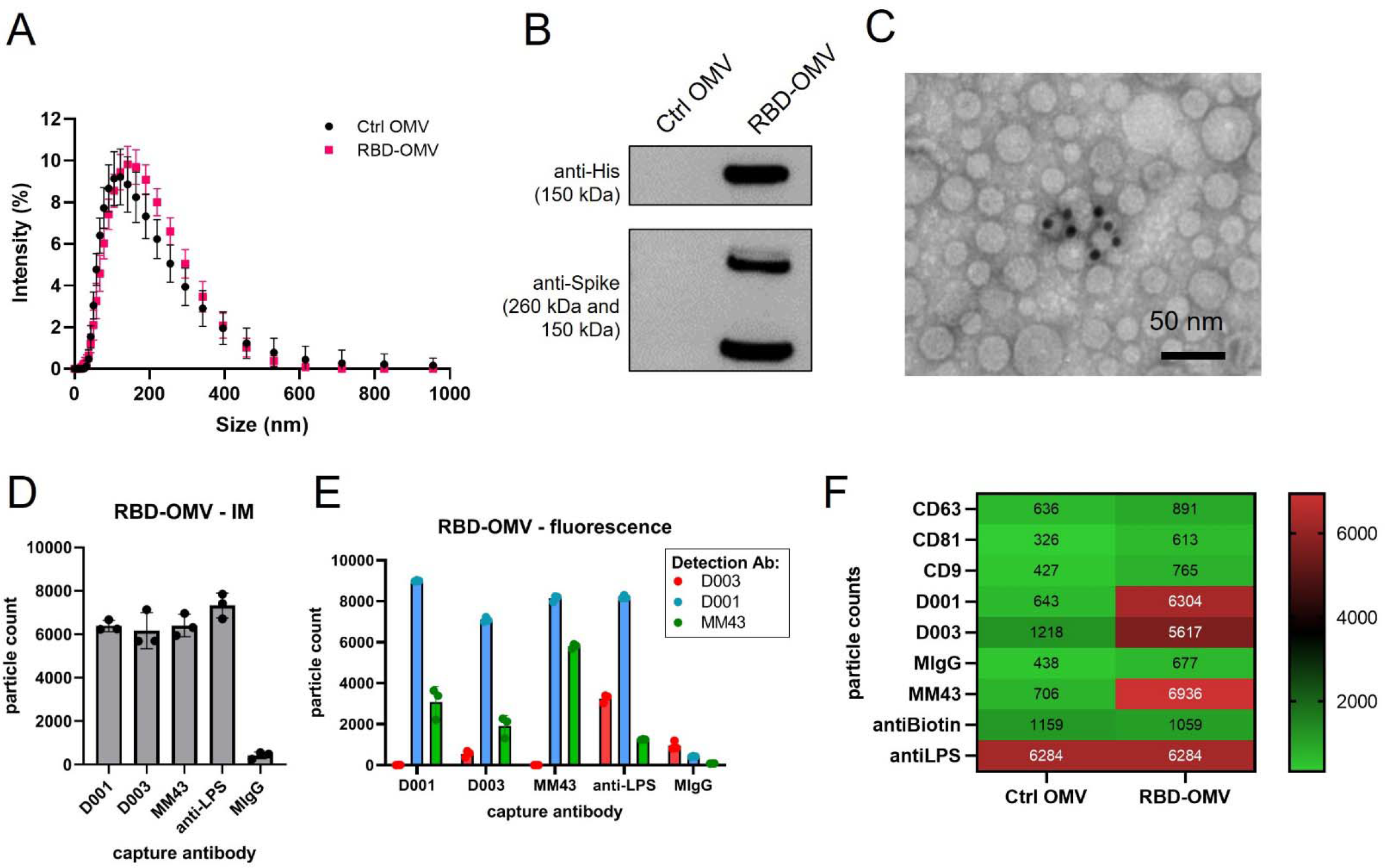
RBD-OMV characterization. A) Particle concentration and size were determined by DLS. Ctrl-OMVs and RBD-OMVs had comparable particle size distribution, with a mean diameter of 118 nm for Ctrl OMV and 125.6 nm for RBD-OMVs. B) Western blot of Ctrl-OMVs and RBD-OMVs probed with anti-His and anti-Spike antibodies. C) Immunogold transmission electron micrograph with anti-Spike-MM43 and streptavidin-gold (10 nm). D) SP-IRIS of RBD-OMVs captured by antibodies against Spike (D001, D003, MM43), anti-LPS, and mouse-IgG isotype control (MIgG). Interferometric imaging (IM) results are light grey bars. Data points show particle counts per capture spot, n=3 capture spots. E) Labeling with fluorescently labeled antibodies D001, D003, and MM43 shows localization of CoV2-Spike epitopes on RBD-OMVs (colored bars). Data points show particle counts per capture spot, n=3 capture spots. F) Heatmap of SP-IRIS data comparing RBD-OMVs from (D) and Ctrl-OMVs. Particle counts for each marker were normalized by LPS content (see also Supplementary Figure S2).

We further characterized the conjugated OMVs by various methods in an attempt to satisfy the recommendations of the minimal information for studies of EVs ^20,21^ (although these guidelines are written mostly for studies of mammalian EVs). Dynamic light scattering (DLS) and Nanoparticle Tracking Analysis (NTA) showed that unconjugated OMVs (Ctrl-OMV) and RBD-OMV are similar in size (Figure 3A and Figure S1D). Immunogold electron microscopy detected RBD on the surface of OMVs (Figure 3C). Multiple factors may influence the accuracy of using immunogold labelling for quantification purpose, including the sample concentration, the accessibility of the epitope to the labeling antibodies, the fixation methods, and incubation time. In addition, in our formulation, antigen may be masked, for example, by steric hindrance by RBD glycans. Thus, the immunogold labeling data presented in Figure 3C should be interpreted as qualitative rather than quantitative. We used SP-IRIS ^22^ to further validate the surface display of RBD on OMVs. This method uses surface-immobilized antibodies to capture nanoparticles, quantify them by interferometric measurement, and subsequently phenotype them using fluorescently labeled antibodies. We used custom chips that were printed with various antibodies against CoV2-Spike (D001, D003, MM43), as well as anti-LPS, which captures all OMVs (Figure 3D). We observed comparable capture of RBD-OMV by the anti-Spike antibodies and anti-LPS by interferometric measurement, consistent with a large percentage of successfully RBD-conjugated OMVs. Furthermore, RBD was detected on the LPS-captured OMVs by fluorescently labeled anti-Spike clones D001, MM43, and, to a lesser extent, D003 (Figure 3E). Ctrl-OMV were captured only by anti-LPS (Figure 3F and Supplementary Figure S2A), and captured particles could not be labeled with fluorescent anti-Spike antibodies (Supplementary Figure S2B).

### Vaccination, virus challenge, and mass/temperature measurements

Next, we evaluated the efficacy of the RBD-OMV vaccine in a recently described SARS-CoV-2 hamster model ^18^ including both biological sexes ^23^. Three groups of 8 hamsters (4 males and 4 females) were inoculated intranasally with Ctrl-OMVs, RBD-OMVs or vehicle on day 0, day 14, and day 28 in a prime-boost-boost regimen (Figure 4A). The animals were challenged with 10^7 infectious units of SARS-CoV-2 on day 44. Body temperature and mass were measured weekly before virus challenge and daily after challenge. No differences in body temperature were measured between the different treatment groups throughout the course of the study (males and females displayed in Figure 4B and C), consistent with previous findings ^18^.

**Figure 4.**
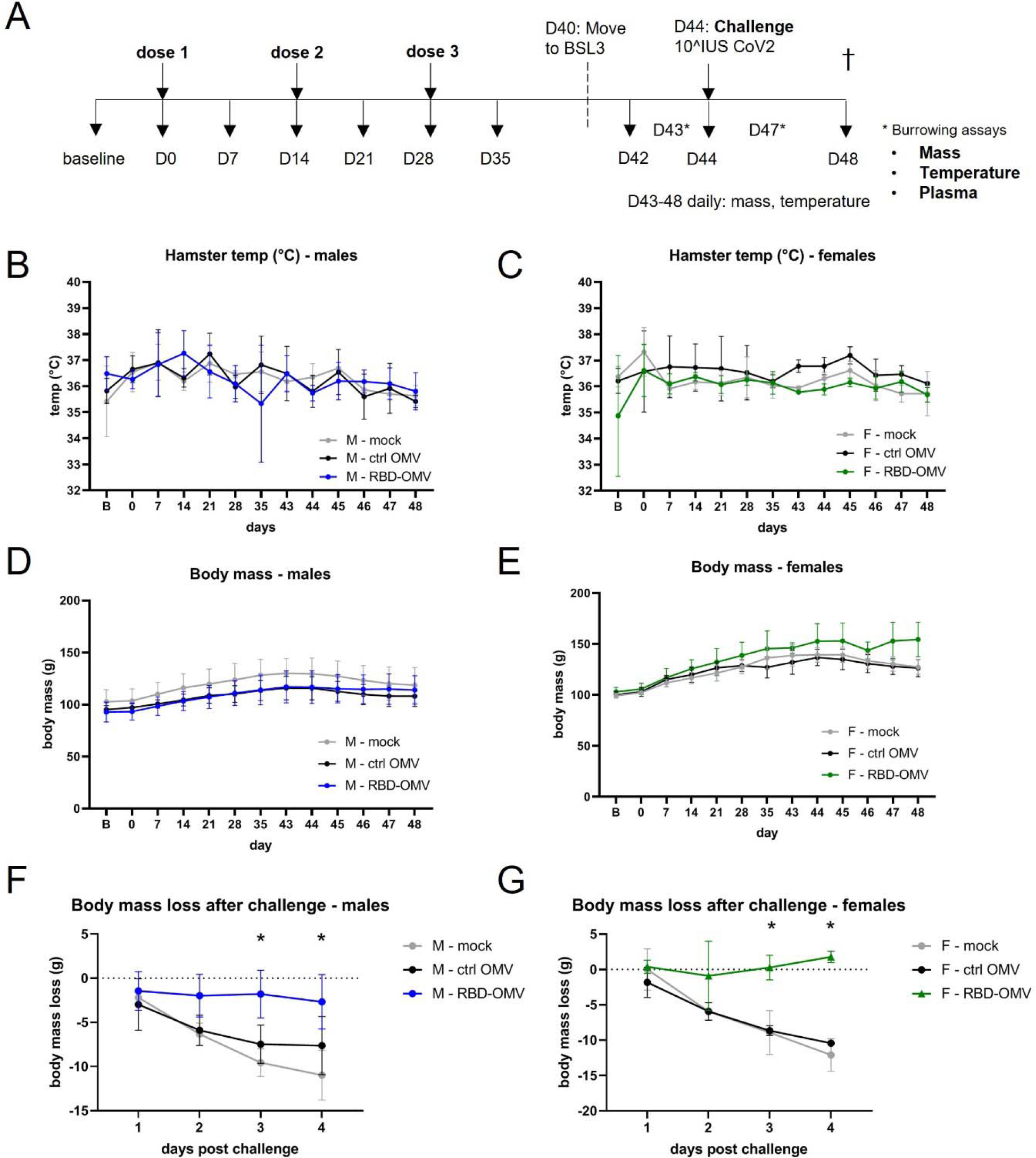
RBD-OMV vaccination prevented loss of body mass after challenge with intranasal SARS-CoV-2, but did not affect body temperature or burrowing behavior. A) Syrian golden hamsters (4 males and 4 females per treatment group) were vaccinated on days 0, 14, and 28 with RBD-OMVs, control OMVs, or mock solution. Hamsters were challenged with 10^7^ infectious units of SARS-CoV-2 on day 44. B-C) Body temperature was monitored *via* a subdermal chip weekly before and daily after virus challenge. D-E) Body mass was monitored weekly before and daily after virus challenge F-G) Mass on days 1-4 post-challenge was measured relative to body mass on day 42. For each day post-challenge, differences in mass loss between groups were tested by one-way ANOVA with Tukey’s post-hoc test, n = 4, * p < 0.05.

### RBD-OMV-vaccinated animals avoided body mass loss after virus challenge

Body mass was previously found to be a reliable indicator of SARS-CoV-2 disease in the model ^18^. Body mass did not differ significantly between the vaccination groups prior to virus challenge (males and females in Figure 4D and E), However, compared with body mass on the day of virus challenge, the body mass of Ctrl-OMV and vehicle groups consistently decreased over four days, reaching significant declines on days 3 and 4 post-challenge. In contrast, RBD-OMV-vaccinated animals avoided this body mass loss, and indeed the vaccinated females had slightly increased average mass by day 4 (Figure 4F and G).

### No significant differences in food burrowing behavior

Food burrowing has been proposed as a surrogate of wellbeing for laboratory rodents including hamsters, in that decreased food burrowing may betray underlying pathology ^24^. We performed burrowing assays at one and three days post-challenge by measuring the amount of food before and after a 24-h interval. There was no difference in burrowing behavior between the groups at one and three days post-challenge (Supplementary Figure S3). There were also no clear differences in burrowing behavior between males and females (Supplementary Figure S3).

### RBD-OMV vaccination elicited RBD-specific plasma IgG responses

Next, we tested whether the RBD-OMV vaccine elicited the production of plasma IgG directed against Spike-RBD. Both males and females in the RBD-OMV group had high plasma IgG titers on day 42, while IgG against Spike-RBD was below the limit of detection in both control groups (Figure 5A). We then examined plasma IgG production longitudinally in the RBD-OMV-treated animals (Figure 5B). After one dose of the vaccine, most animals had detectable Spike-RBD-specific IgG in plasma by day 7, and all by day 14. After the first boost on day 14, IgG levels increased to their maximum levels and were not further increased after the second boost on day 28. Male and female hamsters had comparable IgG titers, with no clear differences in IgG production kinetics.

**Figure 5.**
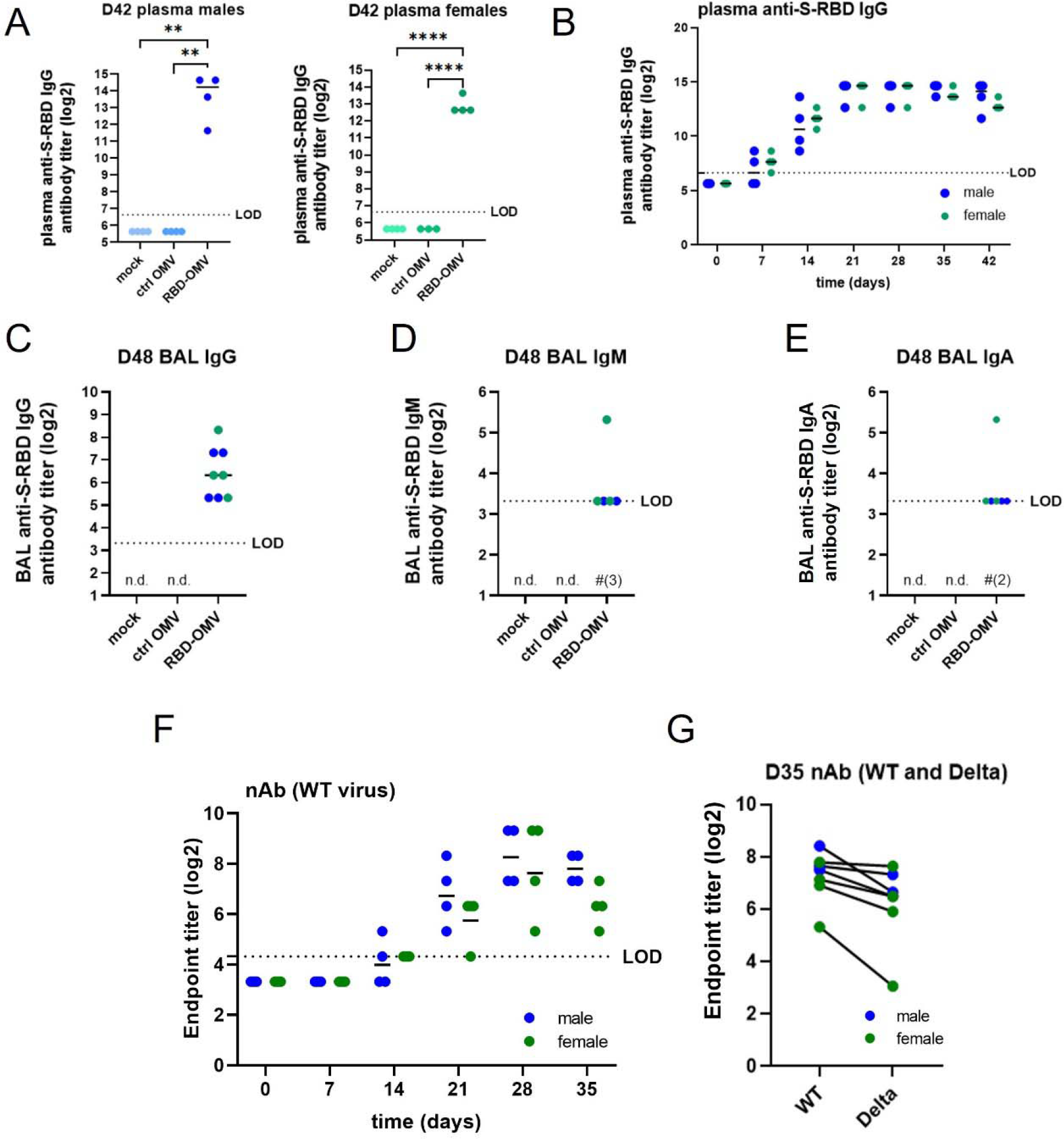
RBD-OMV induced anti-S-RBD-specific IgG in male and female hamsters. A) Pre-challenge anti-S-RBD IgGs was measured by ELISA for day 42 plasma of males and females of all groups. B) anti-S-RBD IgG titers were determined in plasma of RBD-OMV immunized animals collected at different timepoints during the vaccination phase. C) Anti-S-RBD IgG, D) IgM, and E) IgA were determined in day 48 BAL fluid by ELISA. Statistical significance was assessed by one-way ANOVA with Tukey’s post-hoc test, ** p < 0.005, *** p < 0.001, **** p < 0.0001. n.d. = not detected (for all subjects). LOD = limit of detection. Note that in D) and E), levels for most subjects were just above the LOD; in these panels, for RBD-OMV, # is used to indicate the number of subjects in which antibodies were not detected. F) Neutralizing antibody activity against WT virus was measured in plasma of RBD-OMV immunized animals collected at different timepoints during the vaccination phase. G) Neutralizing antibody activity against WT and Delta variants was measured using day 35 plasma. There was no statistically significant difference between neutralizing antibody activities against WT versus Delta, as assessed by paired t-test.

### Bronchoalveolar lavage: anti-RBD IgG, IgA and IgM

Mucosal antibodies provide a first line of defense against airborne pathogens. Therefore, we determined the levels of mucosal antibodies by measuring IgG, IgA, and IgM in bronchoalveolar lavage (BAL) samples collected on day 48 (4 days post-challenge). Anti-S-RBD-specific IgGs were detected in all male and female hamsters treated with RBD-OMVs, but were undetectable in the Ctrl-OMV and mock groups (Figure 5C). IgM antibodies were detected in 2 out of 4 male hamsters and 3 out of 4 female hamsters in the vaccination group (Figure 5D), and IgA antibodies were detected in 3 out of 4 male and 3 out of 4 female hamsters (Figure 5E); however, most of the detected levels of these antibodies were just above the calculated limit of detection.

### Neutralizing antibody activity against WT and Delta SARS-CoV-2

Neutralization assays provide a functional measure of anti-SARS-CoV-2 antibody-mediated immunity. Neutralizing antibodies in hamster plasma samples were tested using a live SARS-CoV-2 microneutralization assay. Neutralization of the WA-1 virus strain (wild type, WT: identical sequence to the RBD-OMV immunogen) increased starting at day 14 after RBD-OMV vaccination, reached a maximum at day 28, and remained high at day 35 (Figure 5F). Day 35 plasma samples were also tested against the Delta variant to assess cross-reactivity. Neutralization activity against Delta was detected for all immunized subjects (Figure 5G). Albeit slightly lower in some of the animals, there was no statistically significant difference between activity against WT versus Delta.

### Infectious virus load in lungs

Virus titers in the lung were determined using BAL fluids and lung tissue at 4 dpi with a TCID50 assay. Virus titers were significantly (100-to 1000-fold) reduced in lung homogenate of RBD-OMV immunized hamsters compared with both control groups, with nearly undetectable infectious virus in the RBD-OMV animals (Figure 6A). RNAscope® ISH was used as a complementary approach for lung tissue TCID50, showing a similar result (Figure 6B). BAL fluids also showed significantly reduced infectious virus in RBD-OMV immunized hamsters at 4 dpi (Figure 6C).

**Figure 6.**
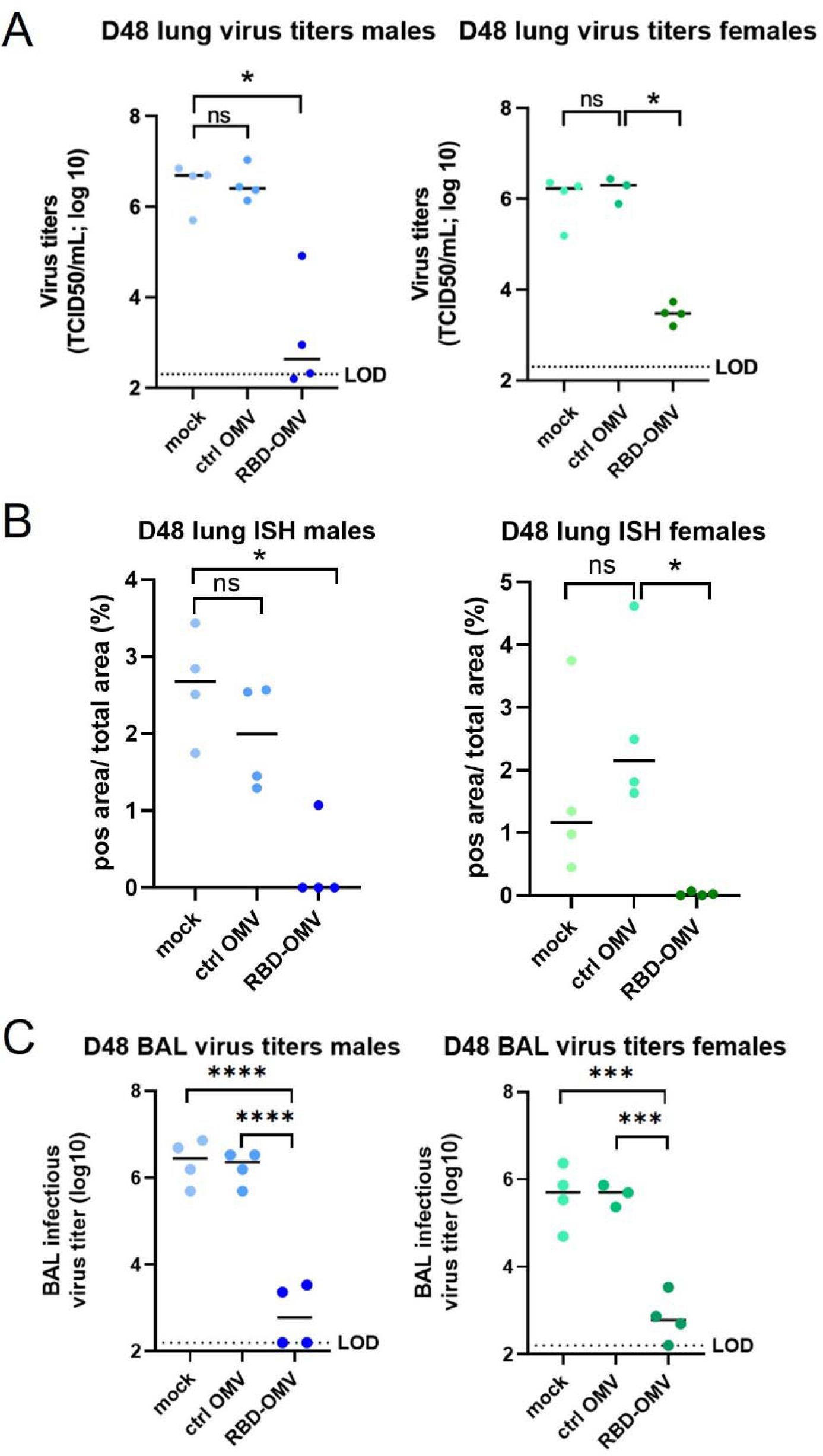
Viral titers in lung. (A) Viral titers in lung tissue measured by qPCR; (B) ISH data of lung tissue; (C) Viral titers in BAL fluid. Statistical significance was assessed by Kruskal-Wallis test, ** p < 0.005, *** p < 0.001, **** p < 0.0001, n=4.

### Gross and histopathologic examination of lungs

At necropsy on day 48, organs were removed and processed as indicated in Methods. Gross examination suggested that the lungs of hamsters immunized with RBD-OMV had fewer focal patches of inflammation and hemorrhagic areas after virus challenge (Figure 7A and Supplementary Figure S4). In addition, the RBD-OMV vaccine group showed less alveolar edema. In contrast, we observed many lesions and inflammation spots in the lungs from the mock and Ctrl-OMV groups. H&E-stained sections of lung were then examined and scored to understand possible differences between the vaccination groups. Lungs of hamsters in the mock (PBS) and Ctrl OMV groups had more focal patches of inflammation, alveolar collapse, and hemorrhagic areas compared with the RBD-OMV vaccinated group (Figure 7B). According to the scoring system, male hamsters vaccinated with RBD-OMV had significantly lower lesion scores than the other groups (Figure 7C). Considering males and females together, the score was also significantly lower.

**Figure 7.**
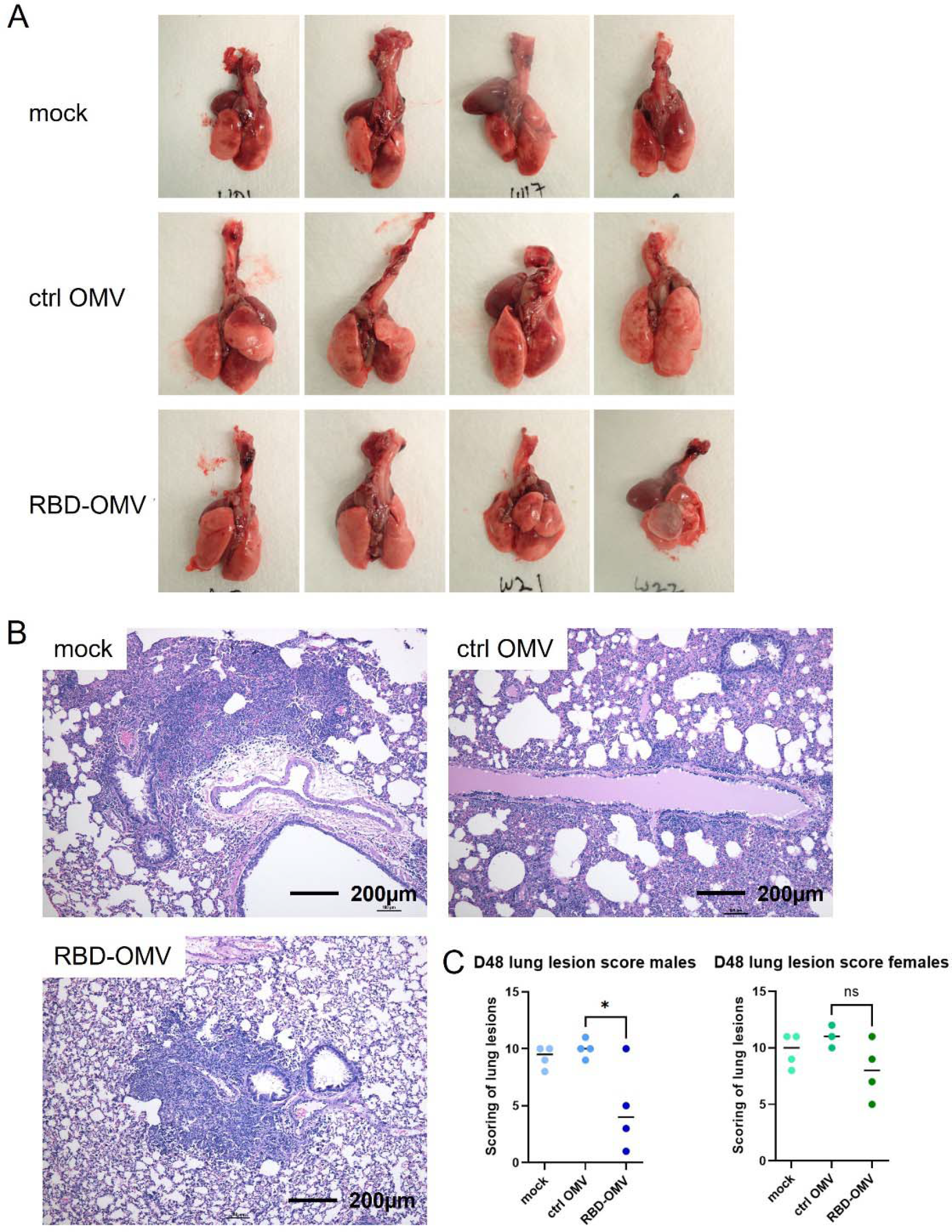
RBD-OMV vaccination reduced pathological lesions in hamster lungs. (A) Gross examination of lungs from hamsters immunized with different formulations (male group). (B) Representative H&E staining of hamster lung sections from each experimental group (20x magnification). (C) Comparison of lesion scores, n=4, * p < 0.05 by one-way ANOVA with Tukey’s post-hoc test.

## DISCUSSION

In this study, we generated and characterized Spike RBD-decorated *S. typhimurium* OMVs and used them to vaccinate hamsters intranasally. RBD-OMVs, but not unconjugated OMVs or a mock vaccination, triggered SARS-CoV-2-specific antibody production as measured in both plasma and bronchoalveolar lavage. Importantly, vaccinated animals had significantly less body mass loss after virus challenge—in some cases, even mass gains—compared with animals in the control groups. Vaccinated animals also had less viral replication and decreased pathological lung lesions. Immunized hamsters showed strong neutralizing antibody titers to the WA-1 challenge virus, which cross-reacted with the Delta variant.

These results demonstrate the feasibility of harnessing OMVs as vaccines, emphasizing several advantages of the platform against SARS-CoV-2 or other viruses. First, scalability: bacteria replicate rapidly, and strains with hypervesiculating properties, like the Salmonella strain used here, produce large amounts of OMVs. Second, versatility: the “plug-and-play” approach allows for decoration of OMVs with a wide variety of antigens or even multiple antigens in the same OMV population. Large batches of OMVs could be prepared, for example, and decorated with appropriate antigens upon emergence of a new viral variant or a new virus. Also, OMV-producing bacteria can be easily engineered and could have their properties “tuned” for specific target groups such as the immunocompromised, elderly, or infants. Third, simplicity of formulation: OMVs are essentially their own adjuvant, obviating the need for adjuvants, which are also sometimes perceived negatively by some in the general public. Fourth, stability: EVs including OMVs are thought to be highly stable, even at room temperature ^25,26^. EVs can also be lyophilized and subsequently stored at 4°C or below ^27^. Of course, stability and efficacy must be tested thoroughly for each specific formulation, but OMV-based vaccines will likely be much easier to store and transport than, *e*.*g*., mRNA vaccines. These properties might recommend OMV vaccines for wider use, especially in geographical areas with limited access to low-temperature refrigeration technologies. Indeed, since our preprint first appeared, we have become aware of two other OMV-based SARS-CoV-2 vaccines in development ^28,29^.

The RBD-OMV vaccine is made with protein produced in mammalian cell culture, which has both advantages and disadvantages. Proteins made by mammalian cells are more likely than those produced by bacteria to have appropriate glycosylation patterns and thus elicit immune responses similar to those that would be expected to real viruses. After treating the recombinant RBD fusion protein with glycosidases, we observed a shift in protein mobility, suggesting that the RBD was indeed glycosylated; however, mass spectrometry is pending and needed to prove the presence of expected glycosylation. As a downside, mammalian cell culture and protein purification are relatively expensive.

There are also potential advantages to the intranasal administration route. As an important barrier against infections, the mucosa are populated by various immune cells, such as dendritic cells, macrophages, T cells, and B cells, which are required to mount an immune response ^30^. An important characteristic of the mucosal adaptive immune response is production of IgA antibodies, which are resistant to degradation in the protease-rich environment of the mucosa ^31^. Intranasal vaccination has been shown to induce IgA in the mucosa ^32^, consistent with our findings. We also found that intranasal vaccination resulted in high IgG levels in plasma, which is supported by previous studies ^32,33^. Thus, intranasal vaccination may optimally result in both mucosal and systemic protection. Intranasal vaccines are also relatively easy to administer, an advantage over existing injectable vaccines. Other studies that used intranasal administration of adenoviral and parainfluenza-based viral vectors against SARS-CoV-2 also reported high neutralizing antibody titers and reduced viral loads in the nose and lungs of hamsters^34,35^, consistent with our findings. Compared with protein subunit/OMV vaccines, viral vectors may elicit stronger immune responses, as they induce sustained antigen expression. However, a known drawback of viral vectors is pre-existing immunity against viral vectors, which is not the case for OMV vaccines. Numerous questions arise from our study. We do not know how different administration routes of OMV vaccines, such as intramuscular, would perform, so future studies might usefully examine this question. We also cannot conclude from the existing data whether or not a single dose of vaccine would have been effective. Blood IgG titers climbed steadily until three weeks after the first inoculation, at which point they plateaued. Since a booster was given at day 14, we do not know if maximum titers would have been reached with just a single dose. The second booster, however, did not appear to have a substantial effect on IgG levels and could likely be omitted in future trials. We also tested only one dose of our vaccine, and we did not compare it with any other vaccine.

We did not observe strong changes in hamster behavior, as measured by the burrowing assay. This assay was developed to measure behavioral dysfunction, for example in severe neurological disorders such as prion disease ^24^. Although SARS-CoV-2 infection may spread to and/or have effects in the human central nervous system ^36^, it is possible that the hamster model does not recapitulate this aspect of COVID-19, or that effects are simply not measurable using the burrowing assay. If this assay is used in the model in the future, it might be revised in some way. For example, hamsters have been reported to prefer burrowing nesting material rather than food ^24^.

An interesting and potentially important finding was the detection of virus in the lungs as well as some possible lung lesions even in the RBD-OMV-vaccinated animals, despite protection against overall disease as indicated by lack of body mass loss. To be sure, real differences between the groups in terms of pulmonary pathology scoring might have been partly obscured by an issue with our study design: the BAL procedure itself may have caused edema and/or bleeding in the lungs of the protected animals, artificially increasing their scores. Other harvested tissues, including but not limited to nasal turbinates and non-lavaged lung, could be examined to help answer this question. We should also note that the challenge dose of the virus far exceeds what is needed for infection, so the vaccine has been subjected to a very stringent challenge. Even so, the possibility that vaccinated individuals could experience some degree of local infection and replication, without disease symptoms, should be considered carefully and might suggest that masking and distancing measures should be continued even by vaccinated individuals until SARS-CoV-2 is eradicated from specific populations.

## CONCLUSIONS

Our work demonstrates that the hamster model is useful for SARS-CoV-2 vaccine studies and that a bacterial OMV-based vaccine platform confers protection against disease in the model. Various advantages of this extracellular vesicle technology render OMVs a possible solution for future vaccine development against SARS-CoV-2 variants, such as the Delta/Omicron variants, as boosters, or for specific populations. OMV-based vaccines also have strong promise for rapid deployment against future emerging infectious diseases.

## MATERIALS AND METHODS

### Molecular cloning of S-RBD constructs

We designed two expression constructs encoding the SARS-CoV2 Receptor Binding Domain (RBD) (isolate Wuhan Hu-1) modified with a flexible linker, a SpyTag motif ^17^ and a 6xHis-tag on the N- or C-terminus, named His-Spy-RBD and RBD-Spy-His, respectively (Figure 1A). Both constructs had a SARS-CoV2 signal peptide (SP) on their N-terminus, and were flanked by a 5’ EcoRI and 3’ BamHI site for cloning. Both constructs were synthesized by IDT (Coralville, IA, USA) and cloned into the pIRESpuro3 vector (cat# 631619, Takara Bio, USA) using EcoRI and BamHI restriction enzymes (R0101S and R0136S respectively, New England Biolabs) and a Rapid Ligation Kit (cat# K1423, Thermo Fisher, USA). The constructs were validated by Sanger sequencing using CMV-F primers (GENEWIZ, South Plainfield, NJ, USA). For expected amino acid sequences, see Supplemental Information 1.

### Recombinant protein production and purification

Expi293F cells (cat# A14527, Thermo Fisher) were maintained in Expi293 medium in vented shaker flasks on a shaker platform maintained at 125 rpm in a humidified 37°C incubator with 8% CO2. Cells were transfected with maxiprep DNA (cat# 12162, Qiagen, Hilden, Germany) of His-Spy-RBD or RBD-Spy-His expression constructs according to the manufacturer’s instructions. Cultures of 3E6 cells/ml were transfected with 1 ug DNA per ml of culture using ExpiFectamine (cat# A14524 Thermo Fisher), and enhancers were added the next day. Six days after transfection, supernatant was harvested, and recombinant RBD protein was purified as follows.

Cell culture medium was centrifuged at 2000 × g for 20 min at 4°C, and the supernatant was collected and filtered through a 0.22-µm Stericup filter. The filtered medium was then incubated with pre-washed Ni-NTA resin (cat# 88222, HisPur™ Ni-NTA Resin, Thermo Fisher) for 2 h on a shaker (∼40 rpm) at RT. Next, the resin-supernatant mixture was centrifuged at 2000 × g for 10 min at 4°C. The supernatant was collected, and the resin was washed with one column volume of wash buffer NPI-20 (buffer composition can be found in the Qiagen Ni-NTA Superflow BioRobot Handbook) four times. Proteins were then eluted off the resin using the elution buffer NPI-250: resin was incubated with elution buffer NPI-250 for 5 min and spun at 890 × g for 5 min at 4°C. Elution was repeated 4 times, and all eluate was pooled into a 50-ml polypropylene conical tube placed on ice. Eluate was concentrated using 10-kDa Amicon Ultra Centrifugal Filters (UFC901096, MilliporeSigma) (for RBD) spun at 2000 × g for 30 min at 4°C or until only 200 to 300 µl remained in the unit. The protein concentrate was washed with phosphate-buffered saline (PBS), stored in PBS/10% glycerol, snap-frozen, and stored at −80°C.

### Production of the OMV-RBD vaccine platform

OMVs were produced from *S. typhimurium* SL3261 ΔtolRA ΔmsbB cells harboring the expression plasmid pHbpD(Δd1)-SpyCatcher as described previously ^16^ and resuspended in PBS. One batch of OMVs carrying Spike RBD was made by adding RBD-Spy-His to OMVs in 7-fold molar excess over the HbpD(Δd1)-SpyCatcher content. A second batch containing an identical amount of OMVs was made by adding Spy-His-RBD in 11-fold molar excess over HbpD(Δd1)-SpyCatcher. Reaction mixtures were incubated for 18 h at 4°C, after which they were pooled. The resulting suspension was diluted with PBS and passed through a 0.45-µm filter to remove potential aggregates. OMV-RBD conjugates were collected by ultracentrifugation (208,000 × g, 75 min, 4°C) and washed by resuspension in PBS containing 550 mM NaCl. OMVs were collected again by ultracentrifugation (293,000 × g, 60 min, 4°C) and resuspended in PBS/15% glycerol. As a control, OMVs incubated with PBS/15% glycerol rather than purified RBD were used. OMV doses were prepared to contain 18 micrograms of total protein, including ∼280 ng of conjugated RBD. Particle count was 3E+10 particles per dose.

### Determination of OMV protein content

OMV total protein content was determined using DC Protein Assay (Bio-Rad). RBD content of OMVs was quantified from Coomassie brilliant blue G250-stained SDS-PAGE gels loaded with bovine serum albumin reference standards. Gels were scanned on a GS-800 calibrated densitometer (Bio-Rad), and the intensities of protein bands were determined using ImageJ (http://imagej.nih.gov/ij/). The content of total HbpD-SpyCatcher-SpyTag-RBD adduct was quantified, after which the RBD content was calculated based on RBD molecular mass.

### Western blotting

OMVs were lysed with 1% Triton supplemented with cOmplete™ Protease Inhibitor Cocktail Tablets (cat# 11697498001, Roche). Samples were mixed with sample buffer with/without dithiothreitol (DTT), heated to 95°C for 10 mins, and subjected to electrophoresis in 4–12% Bis–Tris polyacrylamide gels (Thermo Fisher). Proteins were transferred to Immobilon-FL polyvinylidene difluoride (PVDF) membranes (Merck Millipore), which were subsequently blocked with 5% blotting grade blocker (cat# 170-6404, BioRad) powder in PBS. Blots were probed with primary antibodies: human anti-SARS-CoV-2 Spike (S-ECD/RBD) (cat# bcb03, Thermo Fisher, 1:1000, non-reducing conditions), anti-*Salmonella typhimurium* LPS (cat# ab8274, reducing conditions, 1:1000), and mouse anti-6xHis (ab18184, Abcam, 1:2000, reducing conditions) in 5% blocking buffer in PBS containing 0.1% v/v Tween 20 (PBS-T), incubating overnight at 4°C on a shaker. Blots were washed 3x with PBS-T and incubated for 1 h at room temperature with appropriate secondary antibodies: mouse IgGk-BP-HRP (cat# sc-516102, SantaCruz) or goat anti-human-HRP (cat# 31410, Thermo Scientific), diluted 1:10,000 in 5% blocking buffer. After washing 3x with PBS-T and 2x with PBS, SuperSignal West Pico PLUS Chemiluminescent Substrate (cat# 34580, Pierce) was used for detection with an iBright FL1000 (Thermo Fisher) imager in chemiluminescence mode.

### Immunogold-TEM

Samples (10 µl) were adsorbed to glow-discharged 400 mesh carbon coated ultra-thin grids (Electron Microscopy Sciences 215-412-8400 CF400-CU µL) for 5 min, fixed in 2% paraformaldehyde (EMS, EM grade 16%), briefly rinsed 3x with PBS, and floated on drops for all subsequent steps. All solutions were filtered except for antibodies, which were centrifuged at 13,000× g for 5 min. Grids were placed on 50 mM glycine for 10 min, followed by 3x 2-min rinses in PBS, and exposed to 0.1% saponin in PBS (3 minutes). After a PBS rinse, grids were blocked in 1% BSA in PBS (30 min), followed by incubation with primary antibodies mouse anti-Spike (clone MM43, Sino Biological, 1:100) and mouse anti-*S. typhimurium* LPS (clone 1E6, ab8274, Abcam, 1:200) in 0.1% BSA in PBS (1 h at room temperature). After primary antibody incubation, grids were rinsed in PBS and incubated with streptavidin-gold (10 nm, cat# S9059, Sigma-Aldrich, 1:40) 1 h at room temperature. Grids were rinsed in buffer, followed by a TBS rinse before staining with 2% uranyl acetate (aq.) with Tylose (0.04%) for 30 sec, twice before aspiration. Negative control grids were included in the labeling procedure, leaving out the primary antibody. Grids were dried overnight before imaging the following day on a Hitachi 7600 TEM with XR80 AMT CCD (8-megapixel camera) at 80 kV.

### Single-particle interferometric reflectance imaging sensing (SP-IRIS)

OMVs were pre-diluted 1:500 in PBS, followed by 1:1 dilution in incubation buffer (IB), and incubated at room temperature on ExoView R100 (NanoView Biosciences, Brighton, MA) custom virus chips printed with SARS-CoV2-Spike antibodies (clones D001, D003, MM43, Sino Biological), anti-LPS (1E6, Abcam), and appropriate isotype controls. Chips were processed and read largely as described previously ^37^. After incubation for 16 h, chips were washed with IB 4x for 3 min each under gentle horizontal agitation at 500 rpm. Chips were then incubated for 1 h at RT with fluorescent antibodies against Spike (D001, CF555), (D003, CF647), (MM43, CF488) and LPS (CF647) diluted 1:1000 (final concentration of 500 ng/ml) in a 1:1 mixture of IB and blocking buffer. The chips were subsequently washed once with IB, three times with wash buffer, and once with rinse buffer (all washes 3 min with 500 rpm agitation). Chips were immersed twice in rinse buffer for 5 s and removed at a 45° angle to remove all liquid from the chip. All reagents and antibodies were obtained from NanoView Biosciences. All chips were imaged in the ExoView scanner by interferometric reflectance imaging and fluorescence detection. Data were analyzed using ExoView Analyzer 3.0 software.

### Nanoparticle tracking analysis (NTA)

ZetaView QUATT-NTA Nanoparticle Tracking Video Microscope PMX-420 and BASIC NTA-Nanoparticle Tracking Video Microscope PMX-120 (ParticleMetrix) were used for particle quantification in scatter mode. The system was calibrated with 100 nm polystyrene beads, diluted 1:250,000 before each run. Capture settings were: sensitivity 75, shutter 100, minimum trace length 15. Cell temperature was maintained at 25°C for all measurements. OMV samples were diluted 200,000x in 0.22 µm filtered PBS to a final volume of 1 ml. Samples were measured by scanning 11 positions, recording at 30 frames per second. Between samples, the system was washed with PBS. ZetaView Software 8.5.10 was used to analyze the recorded videos with the following settings: minimum brightness 20, maximum brightness 255, minimum area 5, and maximum area 1000.

### Dynamic light scattering (DLS)

Intensity-based size values of ctrl OMV and RBD-OMV were measured by dynamic light scattering using a Zetasizer Nano-ZS (Malvern Panalytical, UK). Each formulation was diluted 25.5 times in 1x DPBS, and measurements were carried out in 5 replicates using the following settings: manual measurement, 10 runs in replicate, 12 seconds each run, at 25°C.

### Study design, intranasal vaccination and virus challenge, and data/sample collection

All experimental procedures were approved by Johns Hopkins University Animal Care and Use Committee. The program is accredited by AAALAC international. 24 golden Syrian hamsters (*Mesocricetus auratus*, HsdHan®:AURA, 12 females, 12 males, 7-8 weeks old) were purchased from Envigo (Haslett, MI, USA) and were assigned to 3 immunization groups: 1) mock (vehicle) immunization, 2) unconjugated OMV (ctrl-OMV), and 3) RBD-OMV. After 3 days acclimatization, hamsters were weighed and implanted with a subdermal microchip for temperature monitoring and identification. Hamsters were immunized intranasally (10 µl per naris, both nares, using OMV preparations as detailed above) on day 0, day 14, and day 28 under ketamine/xylazine sedation. On days 0, 7, 14, 21, 28, 35, and 42, hamsters were weighed, temperature was measured, and 200-300 µl blood was collected *via* sublingual vein into EDTA tubes. On day 44, hamsters were challenged intranasally with 10^7 TCID50 of SARS-CoV-2 USA/Washington-1/2020, NR-52281 [BEI Resources, virus prepared as described previously^38^] diluted in 100 µl DMEM in an animal biosafety level 3 (ABSL3) facility. Body mass and temperature were monitored daily after infection, up to day 48 (4 days post infection). On day 43 and day 47, food burrowing assays were performed by weighing food before and after a 24 h interval. On day 48, hamsters were euthanized by isoflurane anesthesia followed by blood collection *via* cardiac puncture and bilateral thoracotomy. The right lung lobes were ligated, and bronchoalveolar lavage (BAL) was performed on the left lobe, after which lungs were harvested and placed in neutral buffered formalin (NBF). Trachea, heart, spleen, kidney and liver were harvested and immersed in NBF. Brain was also collected. During the study, one female hamster in the Control-OMV group died for unknown reasons before viral challenge.

### Blood processing

All blood tubes were centrifuged < 1 h after collection for 5 min at 800 × g at room temperature. Plasma was collected from the upper layer and stored at −80°C.

### Serology

Hamster antibody ELISA for RBD-specific IgG, IgA and IgM responses was performed as described previously^38^. ELISA plates (96-well plates, Immunol4HBX, Thermo Fisher) were coated with a 50/50 mixture of His-Spy-RBD and RBD-Spy-His (2 μg/mL, 50 μl/well) in 1X PBS and incubated at 40°C overnight. Coated plates were washed three times with wash buffer (1X PBS + 0.1% Tween-20), blocked with 3% nonfat milk solution in wash buffer, and incubated at room temperature for 1 h. After incubation, blocking buffer was discarded, two-fold serially diluted plasma (starting at 1:100 dilution) or BAL fluids (diluted 1:10) or tissue homogenates (diluted 1:10) were added, and plates were incubated at room temperature for 2 h. After washing plates 3x, HRP-conjugated secondary IgG (1:10000, Abcam, MA, USA), IgA (1:250, Brookwood Biomedical, AL, USA) or IgM (1:250, Brookwood Biomedical, AL, USA) antibodies were added. For IgG ELISA, plates were incubated at room temperature for 1 h; for IgA and IgM ELISA, plates were incubated at 4°C overnight. Sample and antibody dilution were done in 1% nonfat milk solution in wash buffer. Following washing, reactions were developed by adding 100 μl/well of SIGMAFAST OPD (o-phenylenediamine dihydrochloride) (cat# P9187-50SET, MilliporeSigma) solution for 10 min, stopped using 3M hydrochloric acid (HCl), and read at 490 nm wavelength by ELISA plate reader (BioTek 410 Instruments). The endpoint antibody titer was determined by using a cut-off value defined as three times the absorbance of the first dilution of mock (uninfected) animal samples.

### Determination of infectious viral titers

Infectious virus titers in respiratory tissue homogenates were determined by TCID50 assay as previously described ^38^. Briefly, 10% w/v tissue homogenates or BAL fluid were 10-fold serially diluted in infection medium (Dulbecco’s Modified Eagle Medium (DMEM) supplemented with 2.5% fetal bovine serum, 1 mM glutamine, 1 mM sodium pyruvate, and penicillin (100 U/mL) and streptomycin (100 μg/mL) antibiotics), transferred in sextuplicate into 96-well plates containing confluent Vero-E6-TMPRSS2 cells (National Institute of Infectious Diseases, Japan), incubated at 37°C for 4 d, and stained with naphthol blue-black solution for visualization. The infectious virus titers in (TCID50/mL for BAL and TCID50/mg for tissue) were determined by the Reed and Muench method.

### Neutralizing antibody assays

To assess neutralizing antibody titer, SARS-CoV-2/USA-WA1/2020 (BEI Resources) and Delta variant SARS-CoV-2/USA/MD-HP05660/2021 were used. The isolation method for the Delta variant was described previously ^39^. Two-fold serial dilutions of heat-inactivated plasma (starting at a 1:20 dilution) were made in infection medium [Dulbecco’s Modified Eagle Medium (DMEM) supplemented with 2.5% fetal bovine serum, 1 mM glutamine, 1 mM sodium pyruvate, and penicillin (100 U/mL) and streptomycin (100 μg/mL)]. Infectious virus was added to the plasma dilutions at a final concentration of 1 × 10^4^ TCID50/mL (100 TCID50 per 100 μL). The samples were incubated for 1 hour at room temperature, then 100 μL of each dilution was added to 1 well of a 96-well plate of VeroE6-TMPRSS2 cells in sextuplet for 6 hours at 37°C. The inoculums were removed, fresh infectious medium was added, and the plates were incubated at 37°C for 2 days. The cells were fixed by the addition of 150 μL of 4% formaldehyde per well, incubated for at least 4 hours at room temperature, then stained with napthol blue-black. The nAb titer was calculated as the highest serum dilution that eliminated cytopathic effect (CPE) in 50% of the wells.

### Pathology

All tissue samples were immersion-fixed in 10% neutral buffered formalin for at least 7 days under BSL3 conditions. Fixed Specimens were processed routinely to paraffin, sectioned at 5μm, and stained with hematoxylin and eosin (H&E). Pulmonary sections were examined by a pathologist who was blinded to the experimental groups. A subjective score from 1 to 12 was assigned based on the severity of lesions. Semiquantitative lung scoring assessed the degree of involvement, hemorrhage, edema, and inflammation (mononuclear and polymorphonuclear (PMN) leukocytes). Similar scores were obtained on a second review.

### RNA-In Situ Hybridization (RNA-ISH)

SARS-CoV-2 RNA detected by ISH was measured as previously described ^19^. In situ hybridization (ISH) was performed on sections (5 mm thick) of formalin-fixed lung mounted onto charged glass slides using the Leica Bond RX automated system (Leica Biosystems, Richmond, IL). Heat-induced epitope retrieval was conducted by heating slides to 95°C for 15 minutes in EDTA-based ER2 buffer (Leica Biosystems). The SARS-CoV-2 probe (catalog number 848568; Advanced Cell Diagnostics, Newark, CA) was used with the Leica RNAScope 2.5 LS Assay-RED kit and a hematoxylin counterstain (Leica Biosystems). Slides were treated in protease (Advanced Cell Diagnostics) for 15 minutes, and probes were hybridized to RNA for 1 minute. An RNApol2 probe served as a hamster gene control to ensure ISH sensitivity; a probe for the bacterial dap2 gene was used as a negative control ISH probe. For digital image analysis, whole slides containing sections of the entire left lung lobe cut through the long axis were scanned at 20x magnification on the Zeiss Axio Scan.Z1 platform using automatic tissue detection with manual verification. Lung sections were analyzed using QuPath v.0.2.2. For SARS-CoV-2 ISH quantitation, the train pixel classifier tool was used. Within a region of interest (ROI), annotations were created and designated as either positive or ignore, which allowed QuPath to correctly identify areas of positive staining. Percent positive ROI was calculated using positive area detected by the classifier divided by total area of the ROI.

### Data availability

The data that support the findings of this study are available from the corresponding authors upon reasonable request. Source data are provided with this paper.

## ACKNOWLEDGMENT

We thank the National Institute of Infectious Diseases, Japan, for providing VeroE6TMPRSS2 cells and acknowledge the Centers for Disease Control and Prevention, BEI Resources, NIAID, NIH for SARS-related coronavirus 2, isolate USA-WA1/2020, NR-52281. This work was supported by Molecular and Comparative Pathobiology departmental funds (to KWW), the NIH Center of Excellence in Influenza Research and Surveillance (HHSN272201400007C, AP and S.L.K.), and the Johns Hopkins Excellence in Pathogenesis, Immunology Center for SARS-CoV-2 (U54CA260492 S.L.K. and A.P.). KWW and TD are also supported in part by AI144997. The authors thank other members of the Molecular and Comparative Pathobiology Retrovirus Lab for support and helpful comments. Prof. Florian Krammer’s lab (Icahn School of Medicine at Mount Sinai) are acknowledged for providing SARS-CoV-2 protein expression plasmids.

## AUTHOR CONTRIBUTIONS

L.J., W.S.P.J., K.W. J.L. conceived the project. L.J. and T.D. produced the Spy-His-RBD/RBD-Spy-His recombinant protein. W.S.P.J. and H.B.v.d.B.v.S produced the OMV and performed the ligation experiment. M.L. and J.S.V. conducted the hamster study. S.D., R.Z., C.C., K.L., I.S. conducted the immunogenicity evaluation. C.B. conducted the pathology evaluation. L.J, T.D. K.W. wrote this manuscript, all authors critically read and corrected the manuscript.

## Supplementary Information

**Supplemental information 1**

Amino acid sequences of RBD constructs. Signal peptide in yellow, His-tag in magenta, SpyTag in light blue, and RBD in dark blue.

**His-Spy-RBD:**

**MFVFLVLLPLVSSQGSSHHHHHHGSGESGAHIVMVDAYKPTKGSGGTGRVQPTESIVRFPNITNLCPFGEVFNATRFASVYAWNRKRISNCVADYSVLYNSASFSTFKCYGVSPTKLNDLCFTNVYADSFVIRGDEVRQIAPGQTGKIADYNYKLPDDFTGCVIAWNSNNLDSKVGGNYNYLYRLFRKSNLKPFERDISTEIYQAGSTPCNGVEGFNCYFPLQSYGFQPTNGVGYQPYRVVVLSFELLHAPATVCGPKKSTNLVKNKCVNF\*\***

**RBD-Spy-His**

**MFVFLVLLPLVSSQRVQPTESIVRFPNITNLCPFGEVFNATRFASVYAWNRKRISNCVADYSVLYNSASFSTFKCYGVSPTKLNDLCFTNVYADSFVIRGDEVRQIAPGQTGKIADYNYKLPDDFTGCVIAWNSNNLDSKVGGNYNYLYRLFRKSNLKPFERDISTEIYQAGSTPCNGVEGFNCYFPLQSYGFQPTNGVGYQPYRVVVLSFELLHAPATVCGPKKSTNLVKNKCVNFGSGGTGAHIVMVDAYKPTKGSGESGHHHHHH\*\***

**Figure S1.**
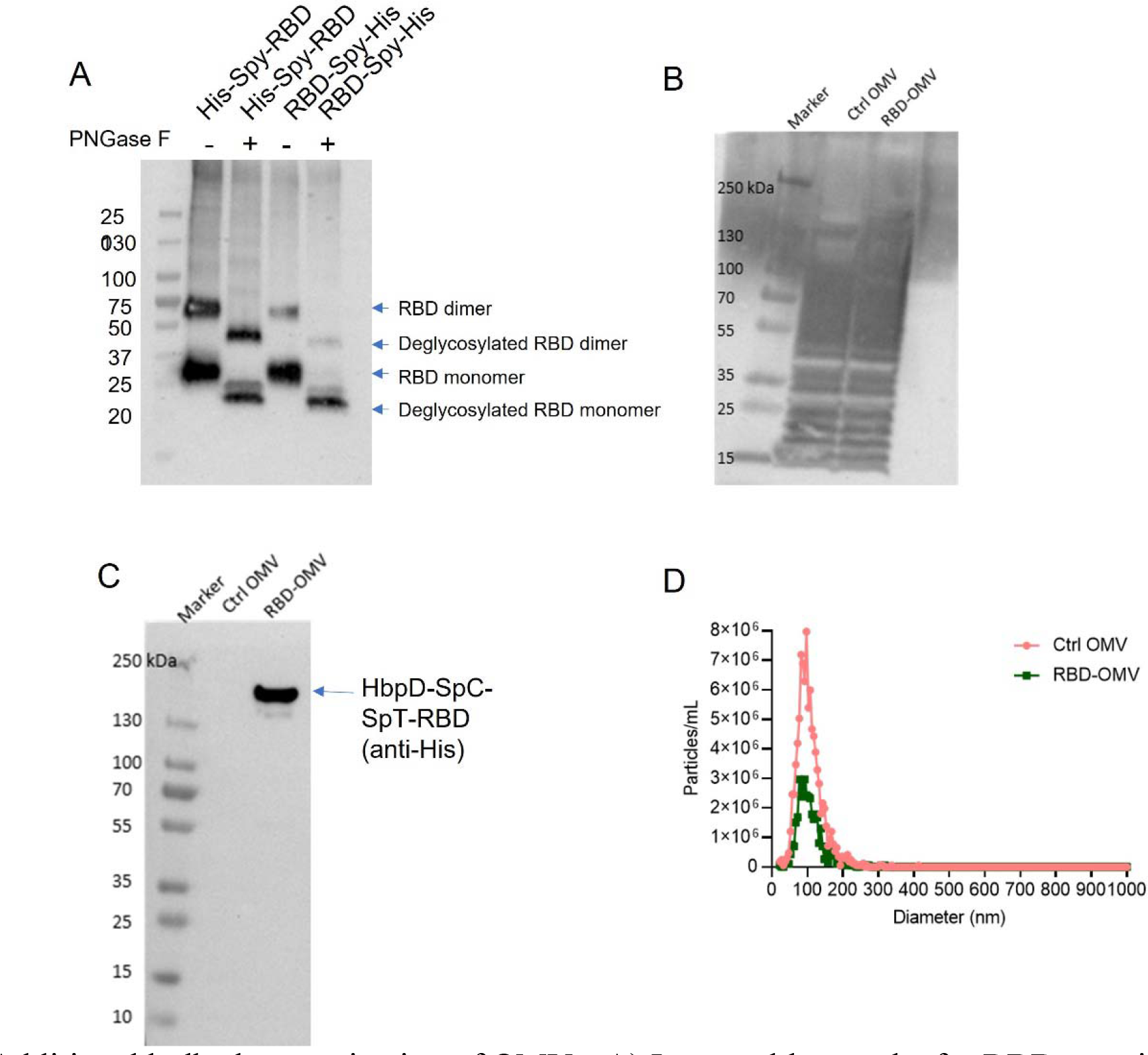
Additional bulk characterization of OMVs. A) Immunoblot results for RBD protein with/without PNGase F treatment; B) Western blot characterization of Ctrl-OMV and RBD-OMV with anti-LPS antibody. C) quantification of RBD in RBD-OMV by anti-His Western blot. D) Characterization of Ctrl-OMV and RBD-OMV by nanoparticle tracking analysis

**Figure S2.**
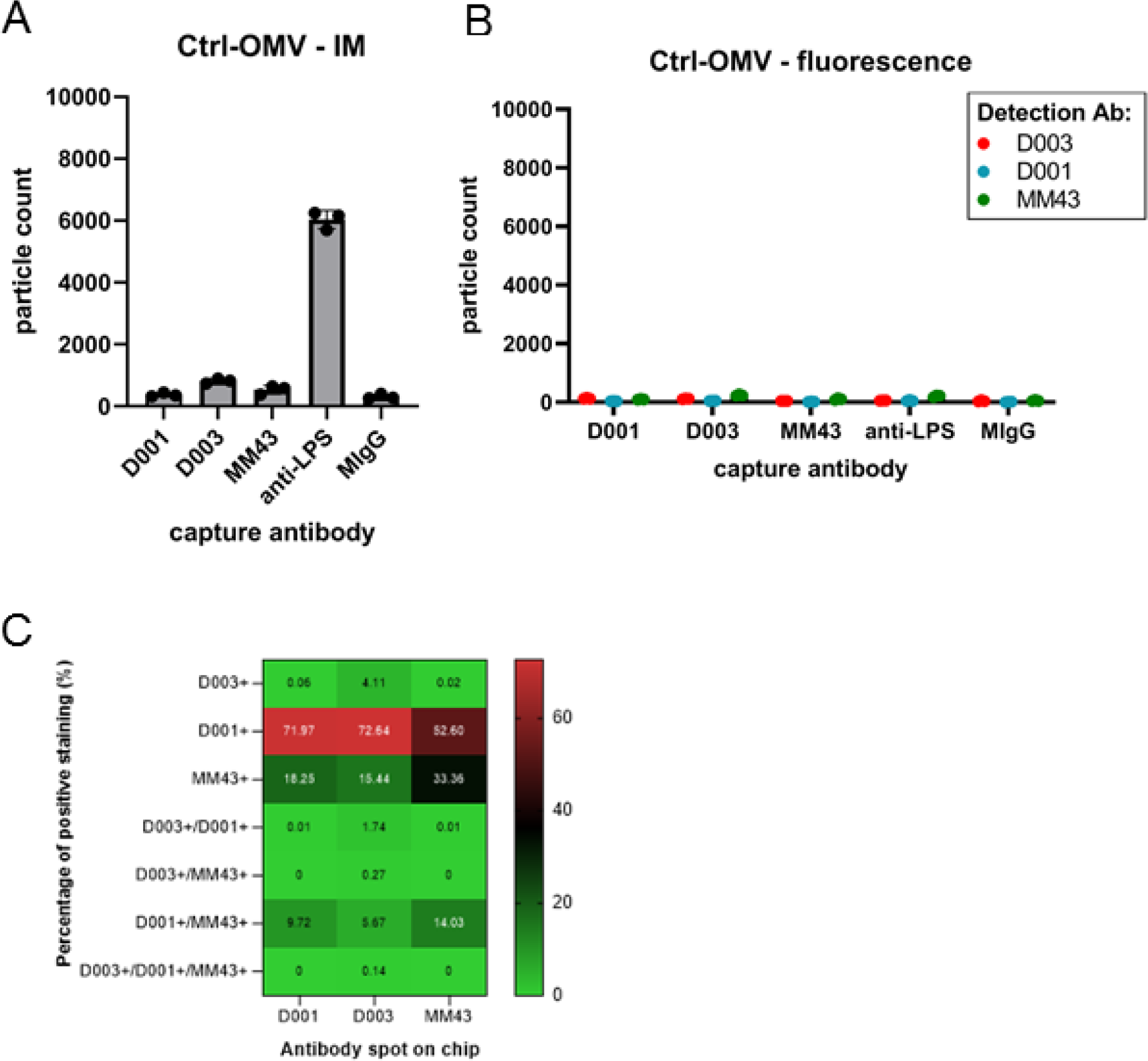
SP-IRIS results for Ctrl-OMVs. (A) Interferometric mode, (B) fluorescence mode. Datapoints show particle counts per capture spot, n=3 capture spots. (C) SP-IRIS results for RBD-OMV, corresponding to Figure 3D-E. Heatmap depicts the percentage of co-localization between fluorescent anti-Spike antibodies on RBD-OMV captured on SP-IRIS chips.

**Figure S3.**
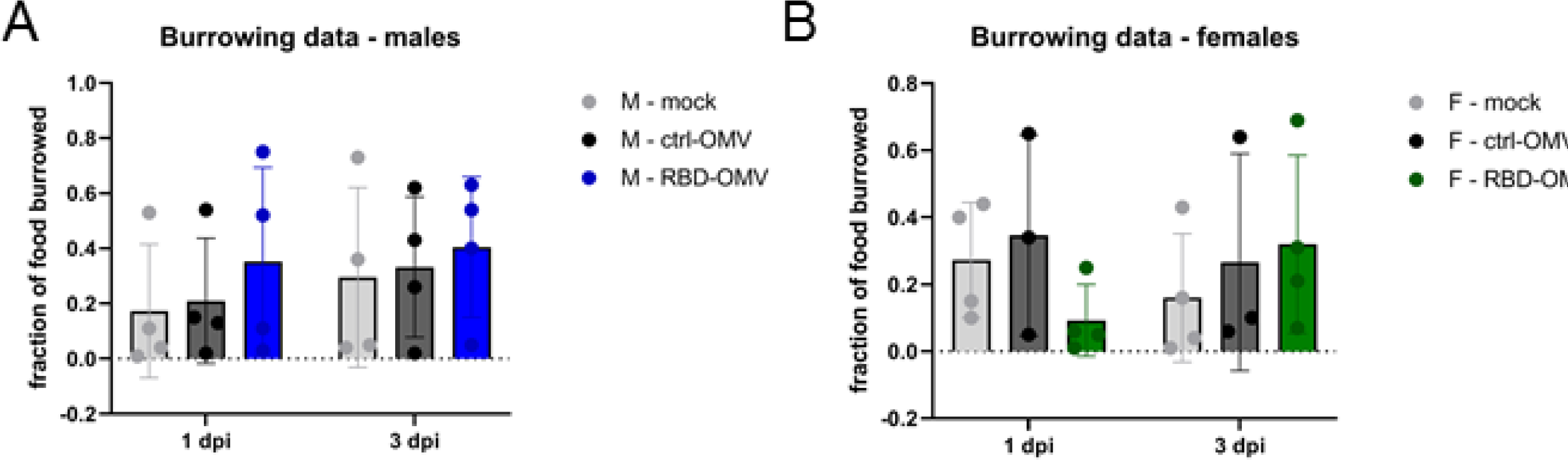
Food burrowing behavior was measured one and three days post-challenge. The fraction of burrowed food was determined by dividing the weight of food after overnight burrowing by the amount of food given to the animals. A) Males and B) females did not show statistical differences in burrowing behavior as analyzed by one-way ANOVA, n=4.

**Figure S4.**
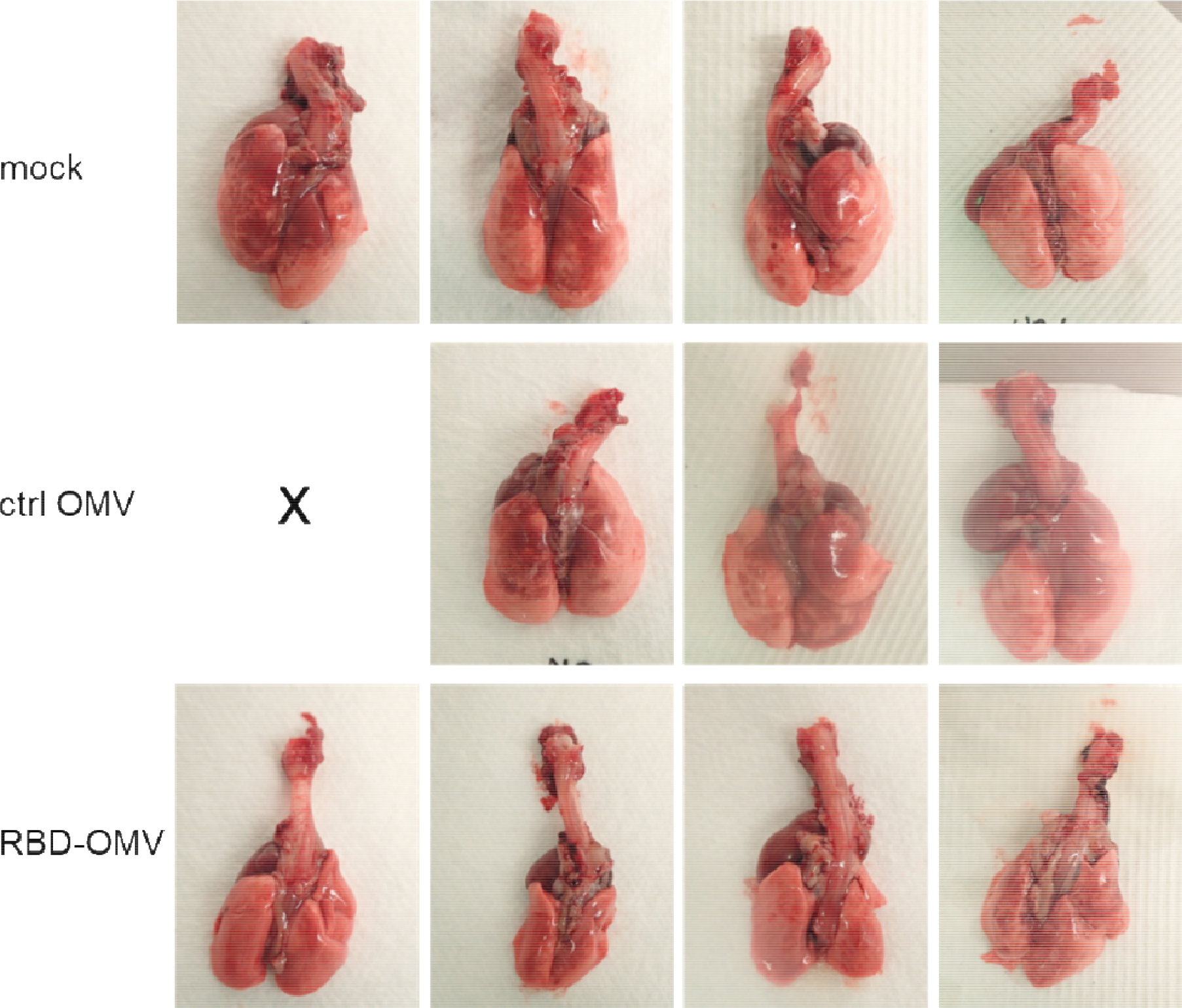
Lungs from female hamsters immunized with different formulations.

